# Viscoelasticity Analysis of Coarse-grained Cytoskeletal Simulations with Cytosim and Cytocalc

**DOI:** 10.1101/2025.10.30.685558

**Authors:** V S Krishna Iyer, Komal Bhattacharyya, Raffaele Mendozza, Peter Sollich, Stefan Klumpp, Yoav G. Pollack

## Abstract

Computational modeling has emerged as a powerful approach to studying cytoskeletal dynamics. The simulation software Cytosim provides intuitive yet flexible simulations of filament polymerization, cross-linking, and motor activity. Here, we present Cytocalc, a lightweight Python toolkit designed to streamline and standardize the analysis of Cytosim simulation output, supporting studies of biological functionality and physical properties of cytoskeletal systems. After introducing Cytocalc and validating it, we use it to establish a new workflow for quantifying network viscoelasticity from Cytosim simulations. Specifically, we determine the complex shear modulus of cross-linked networks and quantify how the storage modulus increases with cross-linker density. The cross-linker dependence of the network’s elasticity exhibits two regimes, a scaling regime consistent with elasticity arising from the suppression of thermal bending fluctuations of filaments as well as a much weaker dependence at high cross-linker concentration.

## 1. Introduction

The cytoskeleton is a dynamic filamentous network that supports essential processes such as cell division, migration, and morphogenesis [1–3]. Its capacity to reorganize and generate forces in response to mechanical cues emerges from the collective behaviour of actin filaments, microtubules, motor proteins, and cross-linkers [4, 5]. Key findings on component dynamics, such as burst-like actin severing by cofilin [6] and rapid microtubule catastrophe–rescue cycles [7] offer important clues about the link between molecular events and cell-scale mechanics. In particular, the cytoskeleton has been shown to underlie the cell’s viscoelastic properties [8], including the characteristic power-law response of living cells [9,10]. Understanding these rheological properties is therefore central to connecting microscopic interactions with emergent mechanical behavior.

Traditionally, such responses have been investigated using simulations of 2D or 3D lattice-based networks [11], or systems with semiflexible filaments represented as bead–spring chains either with [12] or without [13, 14] permanent cross-linkers. These approaches, however, lack the ability to capture dynamic, active features of cytoskeletal networks such as transient cross-linking, filament polymerisation and depolymerisation. Additionally, several theoretical models ranging from polymer theory to mesoscopic models of disordered systems have been proposed to rationalize the physical mechanism leading to the power law rheology exhibited by the cytoskeleton [15, 16], but a complete description is still missing.

Cytosim [17, 18] is an agent-based simulation platform that overcomes these limitations, enabling modelling of various active cytoskeletal components. It has been applied to a wide range of cytoskeleton-related biophysical systems and processes including actomyosin contractility [19], centrosome positioning [20], and actin dynamics in endocytosis [21]. Its open-source code, extensible object model, and efficient Brownian dynamics engine [17] have made it popular with both biologists and physicists. Despite its capabilities, no study has systematically quantified rheology using Cytosim, leaving unaddressed the gap in our understanding of how microscopic interactions give rise to emergent mechanical properties in realistic, dynamic networks.

Furthermore, the analysis pipeline surrounding Cytosim remains fragmented. Users typically rely on ad hoc scripts to parse Cytosim’s plain-text output files and compute quantities such as forces, stresses, or network topology [18] (with the exception of the discontinued cytoslysis package [22]). The problem is further compounded by the variety of output file types of Cytosim, each with potentially different data formats. This lack of standardization complicates post-processing, limits reproducibility, and creates barriers for new users. A unified, well-tested analysis package would eliminate these bottlenecks, streamline Cytosim adoption, and allow researchers to focus on scientific questions rather than data wrangling.

To support both routine and advanced analyses, such a package must accommodate the diverse scientific questions posed by Cytosim users. Biologists may be interested in metrics such as contraction speed [19, 23], network turnover [24], or spatial patterning [25], while physicists often seek to quantify stress relaxation [26–28], defect dynamics [29], or frequency-dependent shear moduli [10, 12, 16]. Despite this diversity, many core observables such as filament spatial distribution, Mean Square Displacement (MSD), and filament–motor interactions are shared across studies [19, 30]. Standardizing how these quantities are extracted and analysed would enable meaningful comparisons between different simulation setups, reduce duplication of effort, and promote reproducible practices. It would also create a common foundation for more specialized extensions tailored to specific research goals.

Here we introduce Cytocalc [31], a lightweight Python package that provides a unified and extensible analysis pipeline for Cytosim simulations. Cytocalc provides support for parsing standard Cytosim output types (termed reports), computing some commonly used observables, and generating basic visualizations suitable for downstream analysis or publication. Cytocalc is an open-source package designed for use in automated analysis workflows and can be extended with new features by contributing users. All code, documentation, and simulation files are freely available [31] to support reproducibility and community use.

Crucially, Cytocalc implements, for the first time, a workflow to systematically measure rheological properties directly from Cytosim simulations. After outlining the implementation and capabilities of Cytocalc and validating its output against published results, we apply it to characterize the rheological properties of various cross-linked actin networks. Using Cytosim to study the rhology of cytoskeletal systems allows to link well-studied macroscopic rheological properties to the molecular properties of their constituents and to the composition of the network. Here we focus on the cross-linker density. We first determine the storage and loss moduli and the crossover with frequency from predominantly elastic to predominantly viscous behavior, for different solvent viscosities. Then, we determine the dependence of the storage modulus on cross-linker density and show that it exhibits two regimes. We find that for relatively low numbers of cross-linkers, the storage modulus scales with numbers of cross-linkers to the third power, indicative of repression of the thermal bending fluctuations of a purely entangled network as the main source of elasticity in agreement with earlier theoretical predictions [13,32]. At higher cross-linker numbers the behavior transitions to a possible power law with an exponent of a half. These results together with the computational workflow proposed here provide a tool for further investigation of cytoskeletal rheology.

## 2. Methods

### 2.1. Core architecture of Cytocalc

Cytocalc follows a simple three-layer conceptual design (Fig. 1): a dedicated parser to read Cytosim report files, an analysis layer, and a lightweight visualisation layer. Clear, well-documented interfaces between these layers allow users to extend or replace components without touching the core logic. The parser supports most standard Cytosim output files (see Appendix A) and is implemented in the `csmparser` module, which handles differences in file structure across report types (see, e.g., Fig. 1d,e). Data from a single simulation time frame is stored in a `CSMFrame` object, while a time series of such `CSMFrame` objects can be collected in a `CSMSimulation` object. These serve as the main entry point for further analysis, where users can access individual frames or call built-in methods to compute time-series or time-averaged observables directly from the `CSMSimulation` object.

**Figure 1:**
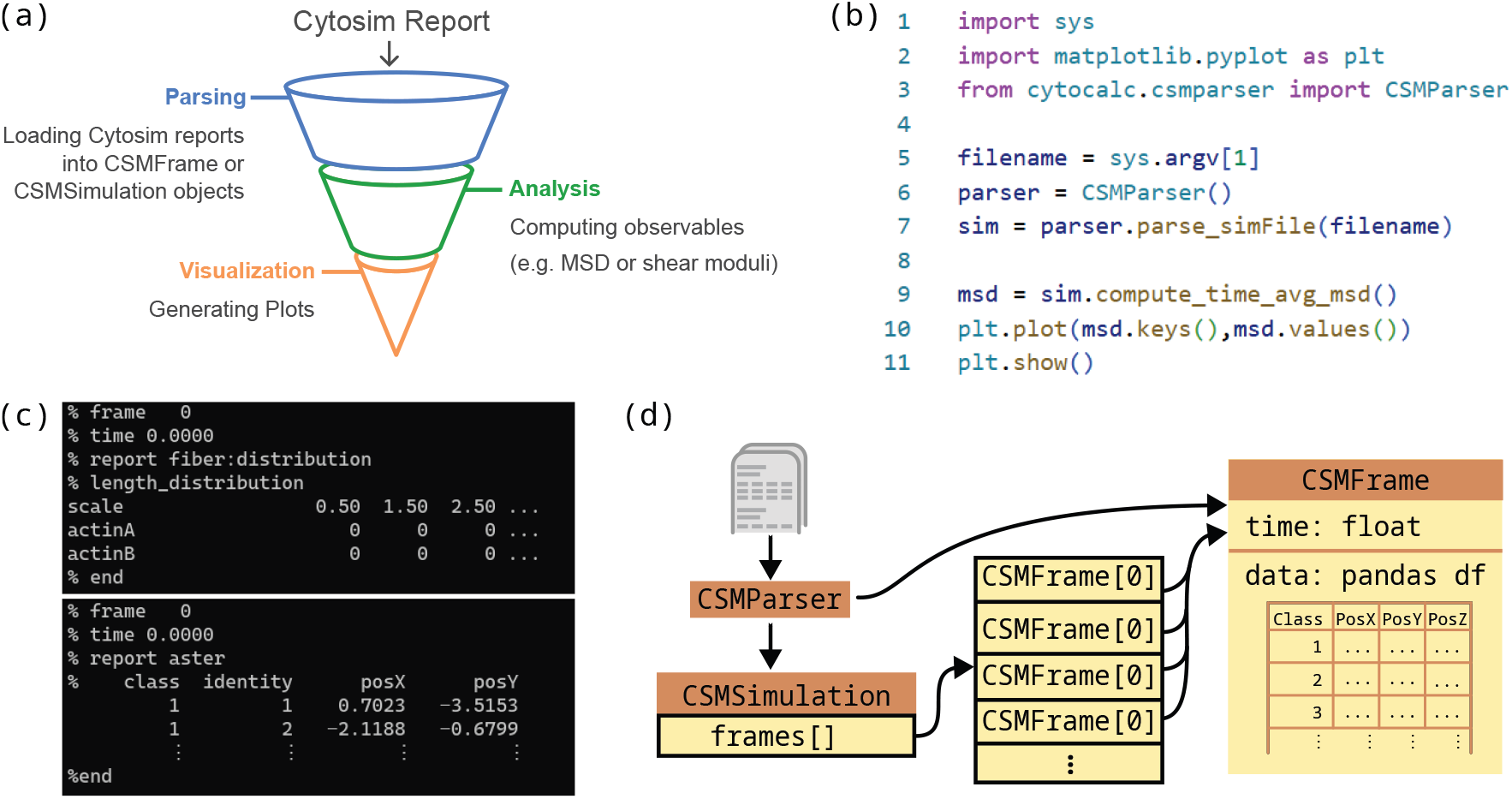
Cytocalc software architecture and workflow. (a) Conceptual three-layer design showing the parser, analysis, and visualization layers. (b) Example minimal script demonstrating the typical workflow from file parsing to Mean Square Displacement analysis and plotting. (c) Sample Cytosim report files illustrating structural differences across output types that the parser handles automatically. (d) Core module architecture showing the relationship between CSMParser, CSMFrame, and CSMSimulation objects in the data processing pipeline.

### 2.2. Extensible analysis layer

The analysis layer provides functions to compute a growing set of observables from the simulation data. The layer includes functions to calculate the spatial distribution of cytoskeleton components and its time evolution, as well as microrheology methods as expanded on in a separate subsection below. Further functions for analysis of non-dynamic filament networks can be easily incorporated from external sources such as freud [33], and future work will add analysis for dynamic networks. A key quantity characterizing filament segment dynamics and the basis of the microrheology measurements described below is the mean square displacement (MSD). Currently, MSD calculations support both displacement-from-origin and time-averaged ensemble methods, with optional correction for periodic boundaries.

### 2.3. Visualization layer for processed data

Cytocalc provides a small set of wrappers built on Matplotlib to simplify the visualisation of key observables such as contraction radius, MSD curves, and viscoelastic spectra. These functions offer sensible defaults for layout and scaling, making them useful for quick inspection or figure preparation. While more advanced or customised plotting is left to the user, saved figures include metadata recording the Cytocalc version used to generate them, supporting reproducibility and basic provenance tracking.

### 2.4. Community-Driven development and maintenance

To facilitate contributions by different research groups and establish Cytocalc as a community-driven tool, the software is released as open-source under the MIT license [31], with the complete source code, scripts and documentation hosted on GitLab. Contributors can add new analysis routines by extending existing classes or introducing standalone modules without modifying the core parser or simulation logic. Cytocalc is written in Python and builds on the standard scientific-Python stack: NumPy, pandas, SciPy, and Matplotlib, interfacing with the freud analysis library [33], to balance ease of installation with computational efficiency. Comprehensive unit tests cover file parsing, object construction, and analysis routines, with automated testing through GitLab Continuous Integration to help maintain stability across future changes. Performance evaluations demonstrate reliable handling of large Cytosim output files ranging from tens of megabytes to several gigabytes under typical desktop conditions.

### 2.5. Microrheology quantification workflow

The complex shear modulus, *G*^∗^(*ω*), characterizes the rheological response of a material to oscillatory shear perturbations of frequency *ω* and infinitesimal amplitude (*linear rheology* ). The real part, *G*^*′*^ (storage modulus), measures the in-phase, elastic-like response, while the imaginary part, *G*^*′′*^ (loss modulus), measures the out-of-phase, viscous-like response. Analogously, it is possible to characterize the material response to an applied infinitesimal shear *stress*, i.e. its deformation compliance, by introducing the complex creep compliance *J*^∗^(*ω*) = 1*/G*^∗^(*ω*).

Cytocalc implements two complementary methods to compute *G*^∗^ and both are employed in this study. These methods rely on the *generalized Stokes-Einstein relations* (GSER), which relate *G*^∗^ to the mean square displacement (MSD) of a probe particle in equilibrium with a continuum bath (see Appendix C for further discussion). These methods have also been applied in cell environments, where they supply insightful though incomplete information [34, 35]. The first method, introduced by Mason et al. [36] relies on an algebraic expansion of the MSD on logarithmic timescales around the studied frequency. This yields an analytical approximation of the GSER, which is accurate for smooth MSD time dependences. The second method, proposed by Evans et al. [37], is a more robust approach valid for viscoelastic fluids. By estimating *G*^∗^ from discrete time measurements of the creep compliance, it avoids artifacts that can result from Fourier-transforms, interpolations and expansions.

## 3. Results

### 3.1. Validation: Actin - myosin network contraction

To ensure the reliability of Cytocalc’s output, we first validated its ability to reproduce known dynamic observables from established Cytosim simulations. As a test case, we considered a published model of actomyosin contraction by Belmonte et al. [19], in which a minimal setup of semiflexible filaments, motors, and cross-linkers generates steady contraction over time (Fig. 2 (a)). Cytocalc computes the radius of the actin network at each time point by extracting filament coordinates from the frame report and evaluating their spatial distribution. The contraction rate is then obtained by a linear fit of the network radius versus time (see details in Appendix B). The resulting contraction profile agrees well with the published results as shown in Fig. 2 (b), where we plot our simulation results together with the theoretical result from Ref. [19] (which also showed good agreement with previous simulations), indicating that Cytocalc reliably extracts dynamic structural properties from Cytosim simulations.

**Figure 2:**
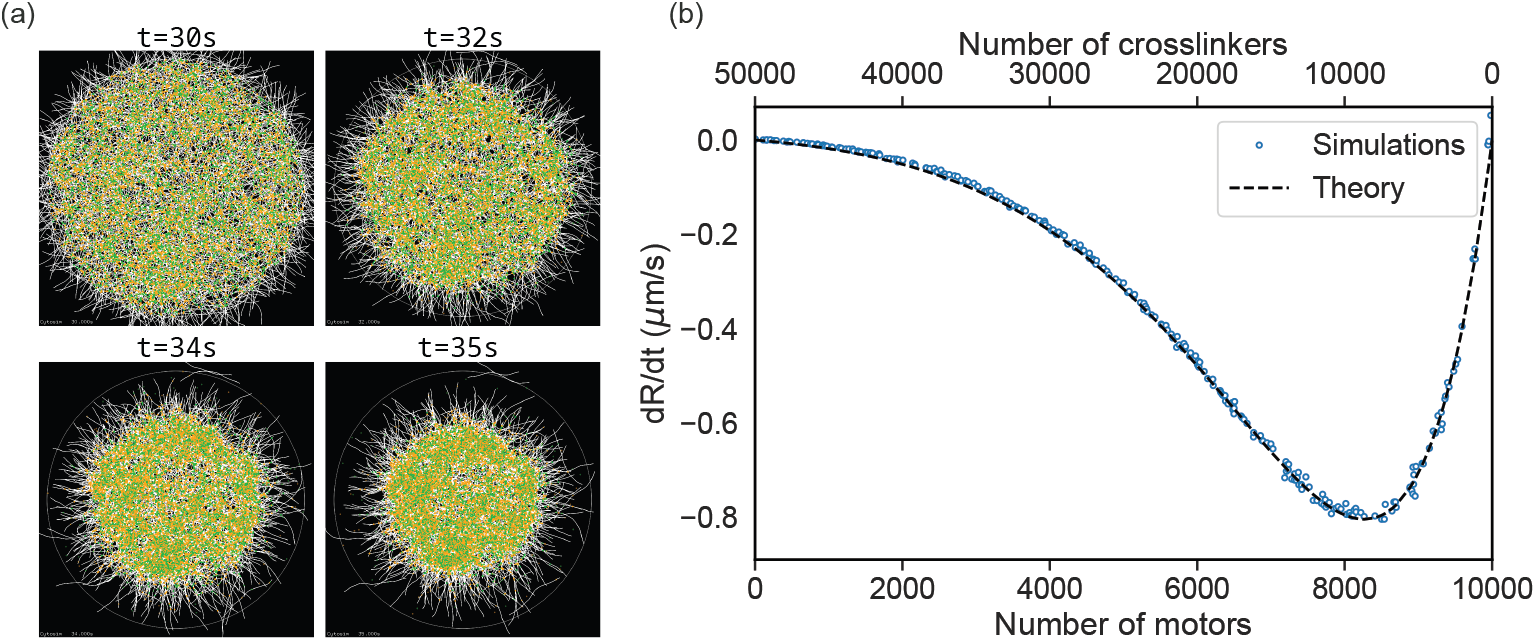
Validation for acto-myosin networks contraction: Reproducing results of Ref. [19]. (a) time series of snapshots of a contracting 2D network (7500 motors) as in Ref. [19]. (b) Rate of change of network radius (*R*) vs number of cross-linkers and number of motors, in good agreement with the theoretical curve from Ref. [19].

### 3.2. Quantification of viscoelasticity of cross-linked actin networks

To evaluate Cytocalc’s reliability also in a soft-matter physics context, where Cytosim has not been utilized so far, we turned to the viscoelastic properties of cross-linked filament networks. These have been previously studied with particle-based simulations (to our knowledge not open source) by Kim et al. [12]. We simulated a dense, 3D, permanently cross-linked, actin-like system (without motors) in Cytosim with 1600 cross-linkers and a solvent viscosity of 0.1 Pa s. The remaining parameters were adapted to those used in the study by Kim et al., see Appendix D, so the comparison with that study provides additional validation of Cytocalc. We tracked the motion of filament segments over time. Cytocalc computes the mean square displacement (MSD) of these segments using a time-averaged ensemble method (Fig. 3 (a)), and infers the frequency dependent complex shear modulus *G*^∗^(*ω*) (Fig. 3 (b)).

**Figure 3:**
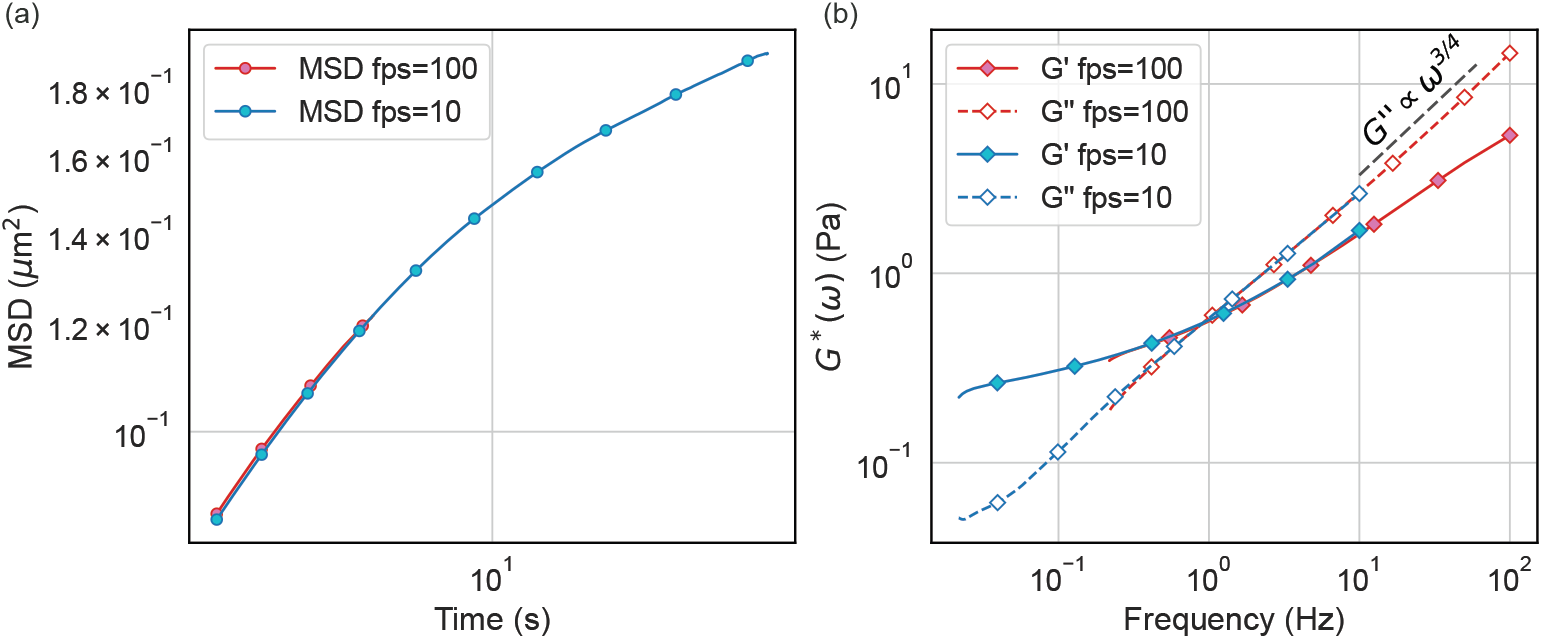
Quantification of frequency-dependent shear modulus from cytosim data. (a) the mean square displacement of filaments over time. (b) the storage and loss moduli calculated from the MSD using Mason’s method [36]. The data points come from two groups of simulations. One with a (post-equilibration) simulation time of 50 seconds and a frame-saving rate (fps: frames per second) of 10 Hz (blue), and the other with a simulation time of 5 seconds and a frame-saving rate of 100 Hz (red). This allows us to sample a wider frequency range without increasing data size. Numbers of cross-linkers 1600, solvent viscosity 0.1 Pa s. See Appendix D for the remaining parameters.

We computed the mean-squared displacement (MSD) of the filament segments (Fig. 3 (a)) and the corresponding real and imaginary component of the complex shear modulus *G*^∗^(*ω*) using the method by Mason et al. [36] (Fig. 3 (b)). To extend the accessible frequency range without increasing the total simulation time by orders of magnitude, we analyzed the same simulation trajectory using two different output frame rates. Specifically, we used data saved at 100 frames per second (pink symbols) and at 10 frames per second (blue symbols). The resulting curves show excellent agreement with each other in the overlapping frequency range, indicating that Cytocalc’s rheology output is robust to the choice of sampling interval. Small deviations near the frequency bounds are expected due to resolution limits. Compared to the study by Kim et al. [12], our approach extends the measurable spectrum toward lower frequencies. This allows us to observe the intersection point of *G*^*′*^ and *G*^*′′*^, below which elasticity dominates, and crossing over to a viscosity dominated regime at higher frequencies, a feature not clearly resolved in the data of Kim et al. *G*^*′′*^ shows a scaling ∼ *ω*^3*/*4^ for frequencies above the crossover point. This can be rationalized by simple scaling arguments of the typical bending mode relaxation following the derivations in Refs. [16] and [38] (see Appendix E), suggesting that cross-linkers hinder the thermal bending of filaments. A further investigation of *G*^∗^ at higher frequencies might be required to confirm this reasoning, which also predicts a scaling of the loss modulus *G*^*′*^ ∼ *ω*^3*/*4^.

Going beyond the previous study [12], we examined how the shape of *G*^∗^(*ω*) depends on the solvent viscosity (shown by the different colors in Fig. 4 (a)). We observe that this parameter influences both the scale of the moduli and their frequency dependence. Increasing the viscosity shifts the crossover from predominantly elastic to predominantly viscous behavior towards smaller frequencies, while both moduli increase. As the solvent viscosity was not explicitly specified in the study of Kim et al. [12], we use it as an effective fitting parameter to align our results with those of their study. The result is shown in Fig. 4 (b): we directly compare there our results to those of Kim et al. [12], setting the solvent viscosity to 110 times that of water, *η* = 110 *η*_*w*_ = 0.11 Pa s. With this choice good agreement is obtained, further validating our approach. These results demonstrate that Cytocalc can be used to calculate microrheological properties of cytoskeletal systems, even extending the accessible dynamic range compared to previous simulations.

**Figure 4:**
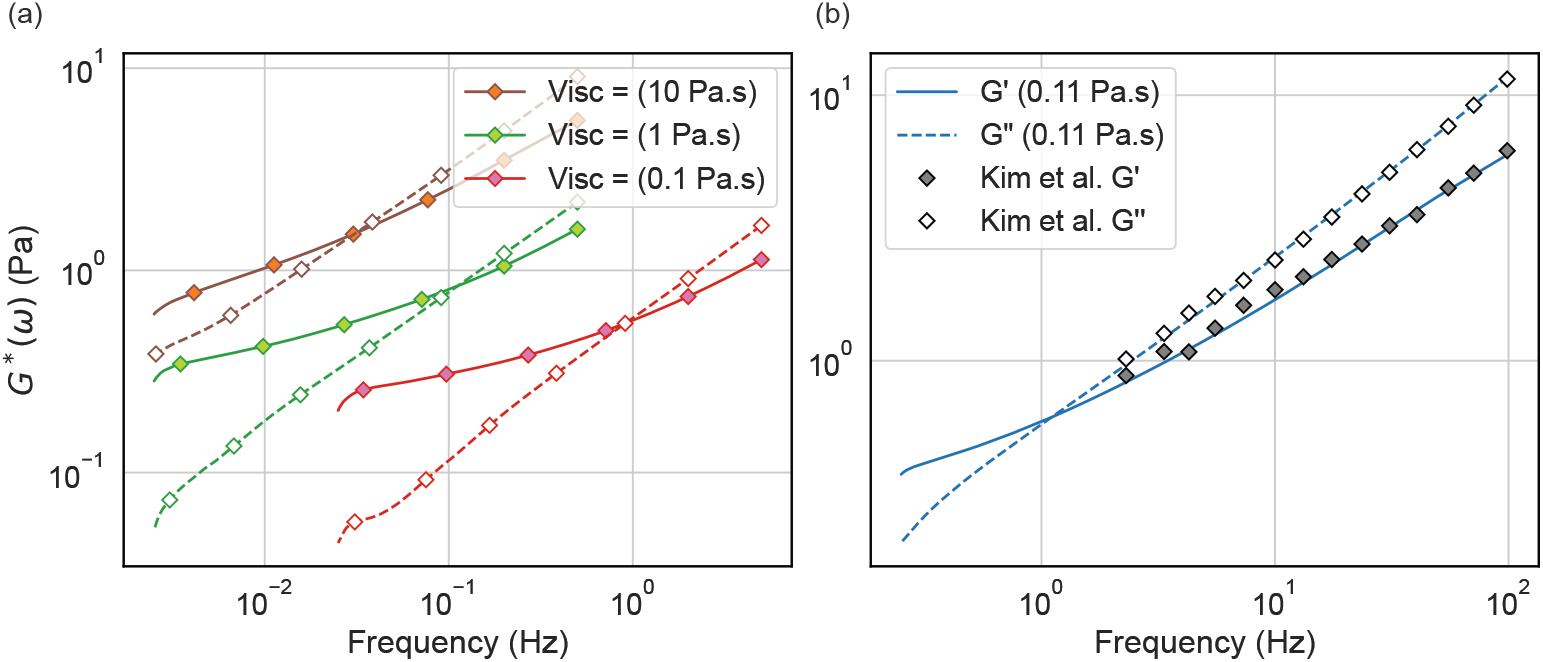
Viscosity dependence of the frequency-dependent shear modulus. Left: The real (lines) and imaginary (dashed lines) components of the frequency dependent shear moduli for different values of medium viscosity: 10 Pa s (orange), 1 Pa s (green) and 0.1 Pa s (blue). Right: Comparison of frequency dependent complex shear modulus for medium viscosity 0.11 Pa s with Kim et al. [12] for the storage and loss moduli. Number of cross-linkers is 1600; see Appendix D the remaining parameters.

### 3.3. Cross-linker–dependence of viscoelasticity

Finally, we use rheology measurements implemented in Cytocalc to study how cross-linker density affects the viscoelastic response of passive actin networks. We simulated motor-free cross-linked networks in Cytosim, systematically varying the numbers of cross-linkers (Appendix D). All other parameters remained unchanged. For each network, we inferred the complex shear modulus *G*^∗^(*ω*) from the MSD. Here, we employed the method proposed by Evans et al. [37] to avoid low-frequency artifacts in *G*^*′*^ in systems with higher cross-linker counts (Appendix C.1).

The storage modulus *G*^*′*^(*ω*) (lines) and loss modulus *G*^*′′*^(*ω*) (dashed lines) are shown for representative cross-linker numbers in the left panel of figure 5 as a function of frequency for representative numbers of cross-linkers, denoted by different colors (data for the full range of cross-linker numbers analyzed appear in Figure D2). As the number of cross-linkers increases, both *G*^*′*^ and *G*^*′′*^ increase across all frequencies, reflecting enhanced stiffness and dissipation. The increase of the storage modulus is, however, more pronounced than the increase in the loss modulus: without cross-linkers, the network is mainly viscous as expected, as expressed by a loss modulus much greater than the storage modulus. With more cross-linkers, the entropy reduction due to entanglement constraints leads to a relative increase in the elastic response, and a crossover appears within our frequency range, similar to Figure 4 and Figure D2. As cross-linker numbers increase even further, the storage modulus surpasses the loss modulus in the entire frequency range we can access, and the material response is effectively elastic.

**Figure 5:**
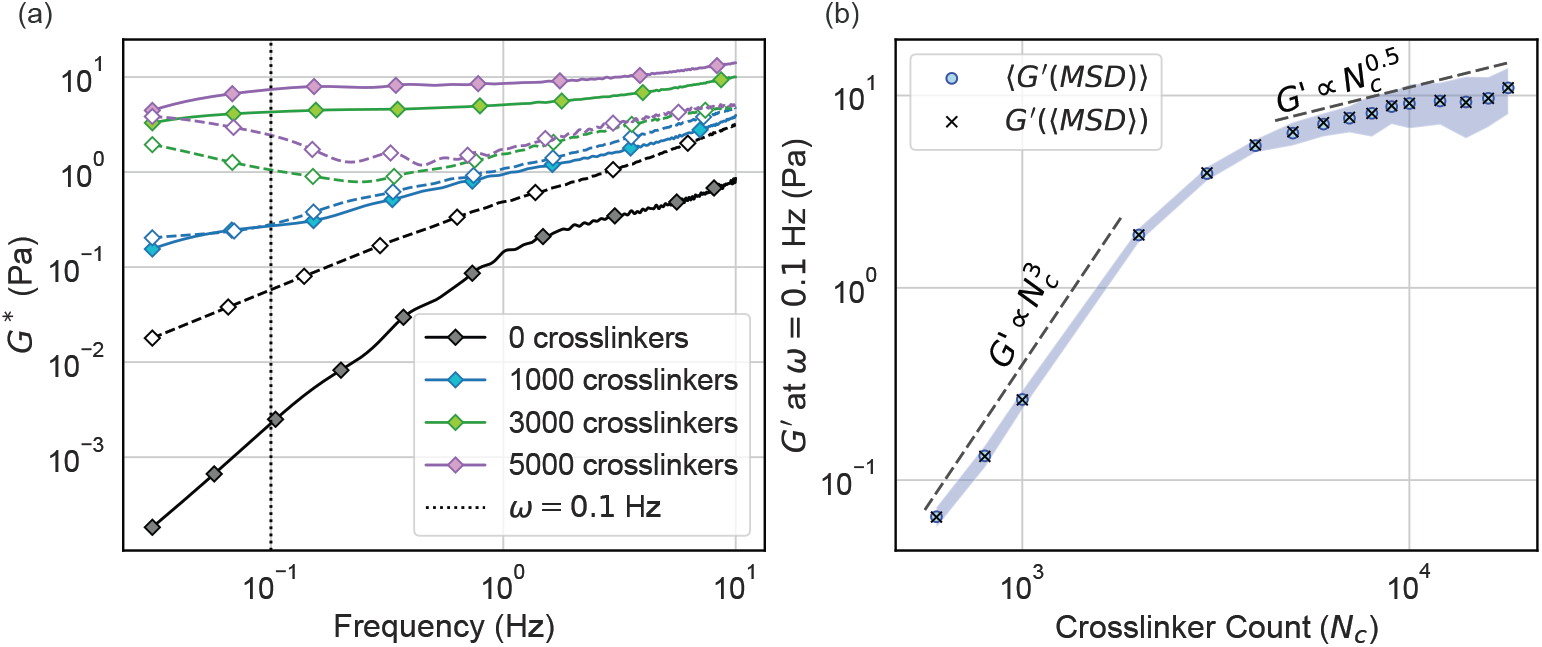
Frequency dependent shear modulus as a function of cross-linker density/ number of cross-linkers. Left: The real (solid lines, filled markers) and imaginary (dashed lines, empty markers) components of the frequency dependent shear moduli, computed using the method proposed by Evans et al. [37] from MSD for different numbers of cross-linkers (shown by the color). Right: Elastic modulus at *ω* = 0.1 Hz against number of cross-linkers. Solvent viscosity 0.1 Pa s; see Appendix D for the remaining parameters.

To assess the effect of permanent cross-linkers on the elasticity of the network we study the scaling of *G*^*′*^ with the number of cross-linkers *N*_*c*_ in the limit of small frequencies as shown in Fig. 5 right. In practice, we determine *G*^*′*^ for a fixed, small, but finite frequency *ω*_0_ = 0.1 Hz, and define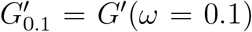. This measure provides a practical proxy for the long-time plateau of the elastic modulus following the approach employed in Ref. [13, 32]. The average of 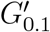 calculated from the MSD average of all realizations (crosses) is consistent with the average taken over the *G*^*′*^ extracted from the MSD of each realization separately (blue circles, with the shaded region indicating the standard deviation), as expected from an equilibrium system.

We observe two distinct regimes in Fig. 5 right: increasing *N*_*c*_, we find a cubic scaling region 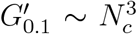, eventually crossing over to an approximate scaling 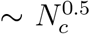 for high cross-linker number. The first region suggests that the low-frequency elastic response is dominated by entropic bending fluctuations of filament segments between adjacent cross-links, as one can understand as follows: if the cross-linkers are approximately uniformly distributed, the average distance between them along a filament is 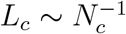.

Each filament segment of length *L*_*c*_ behaves as an entropic spring, with stiffness set by the suppression of thermal bending modes leading to a prediction of 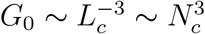, in excellent agreement with our data and consistent with theoretical predictions for affine cross-linked networks of semiflexible polymers [13, 32].

At higher cross-linker densities, however, the dependence of 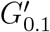 on *N*_*C*_ weakens, approaching an effective exponent close to 0.5. This deviation from the simple affine thermal model suggests that additional mechanisms become important once the network is densely cross-linked. Possible contributing factors include the suppression of nonaffine bending modes, changes in network architecture [39], or the onset of stretching-dominated elasticity [40].

Overall, our results align with current understanding of the mechanics of crosslinked semiflexible polymer networks and demonstrate how Cytocalc can be used to probe scaling behavior in cytoskeletal assemblies.

## 4. Discussion

In this work, we introduced Cytocalc, an open-source Python package for analysing Cytosim simulations. While broadly applicable to a range of problems in the biology and physics of the cytoskeleton, leveraging the versatility of Cytosim as a simulation platform [17], Cytocalc’s initial functionalities are particularly well-suited for investigating the rheology of cytoskeleton networks. We validated the software against established results in cytoskeletal mechanics, including contraction dynamics [19] and linear viscoelasticity [12].

Using Cytocalc, we then determined the frequency-dependent shear modulus of crosslinked semi-flexible filaments. The calculated storage and loss moduli, determined via the method of Mason et al. [37] agree with previous studies [12] and extend the accessible frequency range. There is a characteristic crossover frequency (depending on medium viscosity), where the network transitions from predominantly elastic to predominantly viscous behavior. We observed that for relatively low concentrations of cross-linkers (but still high enough not to constitute a purely entangled melt), the high frequency shear modulus is proportional to (*iω*)^3*/*4^, suggesting an affine response of the cross-linked network [13]. Increasing further the number of cross-linkers, we studied how the low-frequency elastic modulus varies with the cross-linker density and observed a crossover from a thermal affine response 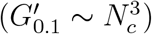 to an approximate scaling 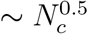, whose physical origin remains an open question for future work. Indeed the integration of Cytosim with the Cytocalc analysis pipeline now enables a streamlined workflow for future studies of cytoskeleton rheology, including for composite networks and uniquely active processes such as dynamic cross-linkers, filament polymerization and turnover.

Cytocalc fills a gap in the Cytosim analysis pipeline by providing a transparent and extensible framework for computing dynamical, structural and mechanical observables from raw simulation output. Its modular design—separating parsing, analysis, and visualization—enables reproducible workflows and lowers the barrier to systematic post-processing. We note a partial functionality overlap with PyCytosim [41], a C-Python interface developed by Serge Dmitrieff that operates on Cytosim source code directly and thus provides comprehensive access to all native Cytosim functions and objects. This offers greater functionality but requires more technical expertise and is less readily integrated into existing research workflows.

While the current Cytocalc version supports a limited set of observables, the codebase is designed for expansion. Additional metrics—such as filament curvature, orientation correlations, or turnover rates—can be incorporated without modifying the parser or core data structures. Looking forward, we envision community-driven development of specialized modules, as well as integration with experimental data streams or other simulation platforms. Indeed, as a modern open-source tool built on standard Python libraries, Cytocalc provides a foundation for building robust, interoperable workflows in cytoskeletal simulation research.

## Acknowledgments

We would like to thank Francois Nedelec, Serge Dmitrieff, Kim Taeyoon, Anastasiia Smagliuk, and Denis Wittig for discussions, feedback and suggestions. We acknowledge support from the Deutsche Forschungsgemeinschaft (DFG, German Research Foundation) – Project-ID 449750155 – RTG 2756, projects A3 (to SK) and A5 (to PKS). Simulations were run on the GoeGrid cluster at the University of Göttingen, which is supported by the Deutsche Forschungsgemeinschaft (Project IDs 436382789; 493420525) and MWK Niedersachsen (grant no. 45-10-19-F-02).

## Declaration of interests

The authors declare no competing interests.

## Data availability statement

The Cytocalc source code can be found on Gitlab [31]. The Cytosim configuration file for the contraction simulation can be found in the SI [42] as well as in Ref. [19]. The Cytosim configuration file for the rheology simulation is found in the SI.

## Appendix A. Supported Report Types

As mentioned in Section 2.1, Cytocalc supports the vast majority of Cytosim’s different output types. To address the variability in the structure of report files, Cytocalc categorizes them into types I, II or III based on the position of the column headers relative to the line declaring the report type. In type I report files, the header follows the report type, whereas type II files contain a comment between the two lines. Type III files have a header column instead of a header row. Table A1 provides a list of supported report file types, classified into the respective categories.

**Table A1:**
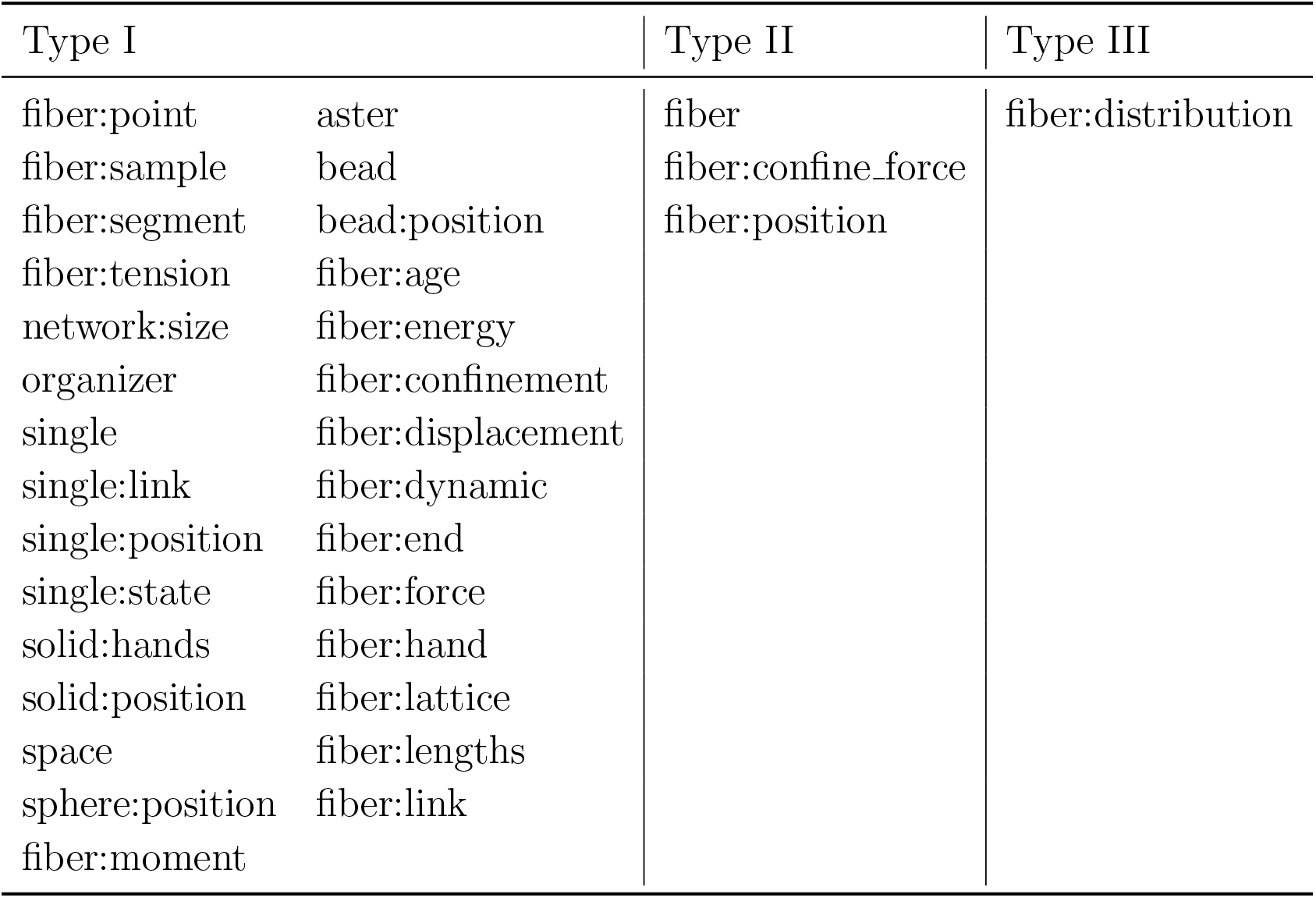
Supported Report Types.

| Type I |  | Type II | Type III |
| --- | --- | --- | --- |
| fiber:point | aster | fiber | fiber:distribution |
| fiber:sample | bead | fiber:confine_force |  |
| fiber:segment | bead:position | fiber:position |  |
| fiber:tension | fiber:age |  |  |
| network:size | fiber:energy |  |  |
| organizer | fiber:confinement |  |  |
| single | fiber:displacement |  |  |
| single:link | fiber:dynamic |  |  |
| single:position | fiber:end |  |  |
| single:state | fiber:force |  |  |
| solid:hands | fiber:hand |  |  |
| solid:position | fiber:lattice |  |  |
| space | fiber:lengths |  |  |
| sphere:position | fiber:link |  |  |
| fiber:moment |  |  |  |

## Appendix B. Extraction of the Contraction Rate from Simulations

In order to measure the contraction rates discussed in 3.1, we simulate a 2D network with motors and cross-linkers using parameters from Belmonte et al. [19]. Following this reference, for each simulation, the network radius was measured as a function of time:

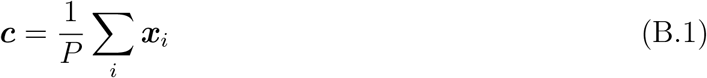

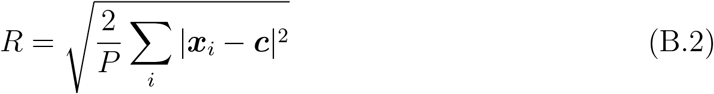

where ***c*** is the center of mass, ***x***_*i*_ are the positions of all the filament segments and *P* is the number of such segments. Eq. B.2 provides an estimate for the radius of a disk given *P* points uniformly distributed on its surface.

**Figure B1:**
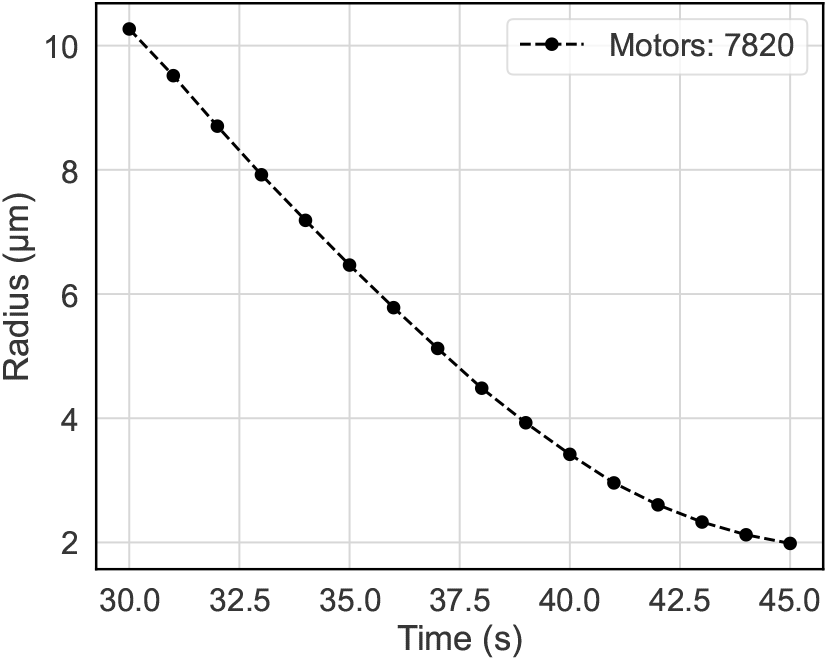
Network radius vs time for an example simulation with 7820 motors and 10900 cross-linkers. The contraction rate *dR/dt* is −0.792 *µ*m/s.

In the work of Belmonte et al. [19], the simulation time was short enough to only capture the regime of contraction linear in time, each run simulating only 5 seconds after the initial equilibration. Running the simulation for longer, we observe that the stochastic contraction mechanism exhausts itself, and eventually the decrease in the network radius saturates. We limit ourselves to the initial linear regime for calculating the contraction rate.

To characterize the contraction rate as a function of motor count, we follow the example of Belmonte et al. and run 300 simulations with motor counts chosen from a uniform distribution between 0 and 10000. In each simulation, the cross-linker count is set to 50000 − 5*N*_*m*_ where *N*_*m*_ is the number of motors in the system. The results are shown in Fig. 2 and discussed in the main text.

## Appendix C. Methods for obtaining the shear modulus from MSD

The mean squared displacement of a particle of radius *a* in a purely viscous fluid is given by:

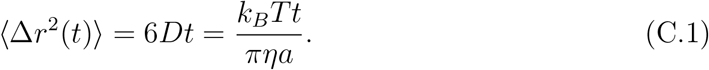

While the MSD, ⟨Δ*r*^2^⟩, is typically described as a function of time, we can, in principle, apply a unilateral Fourier transform [43] to Eqn. C.1 to obtain a frequency dependent mean-squared displacement

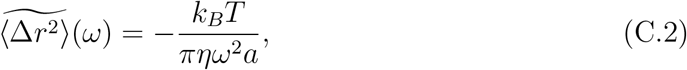

where 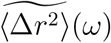 is the MSD in Fourier space.

The Generalized Stokes-Einstein Relation (GSER) similarly relates the 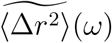 of a particle in a non-Newtonian liquid to a frequency dependent viscosity *η*^∗^(*ω*):

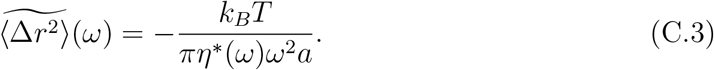

From Eq. C.3 and Eq. C.2, it is evident that the standard Stokes-Einstein relation is retrieved from the GSER for a medium with frequency-independent viscosity *η*. The introduction of *η*^∗^ is a heuristic argument, based on the assumptions that the probe particle is in equilibrium with a continuum bath and that the flow field around the particle does not deviate from Stokes law, independently of frequency. It is then reasonable to assume that the particle diffusion spectrum will be influenced by the viscoelastic spectrum of the bath [36, 44]. The complex shear modulus can then be calculated from the complex viscosity *η*^∗^ as:

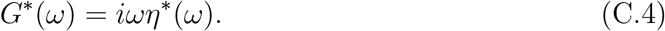

Substituting Eq. C.4 into Eq. C.3 and rearranging gives explicitly:

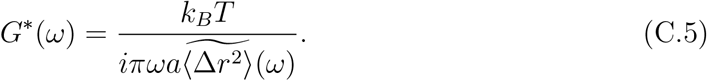

With Eq. C.5 one then has a theoretical framework for extracting the complex shear modulus *G*^∗^(*ω*) from the Fourier transform of the mean squared displacement.

### Appendix C.1. Mason’s method

Using the GSER to calculate the shear moduli requires performing a unilateral Fourier transform on an arbitrary MSD obtained from simulation (or indeed experiment). Such numerical integration can often induce hard-to-control errors. The method proposed by Mason et al. in Ref. [43] is based on a linear expansion of ln (⟨Δ*r*^2^(*t*)⟩) around *t* = 1*/ω*, where *ω* is the frequency being considered. After performing the Fourier transform one obtains

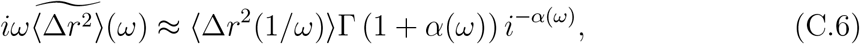

where Γ(1 + *α*) is the Gamma function and *α*(*ω*) is defined as:

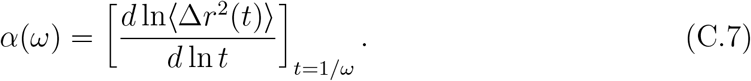

Using this approximation in Eq. C.5 one finds the following expressions for the storage (*G*^*′*^) and loss (*G*^*′′*^) moduli:

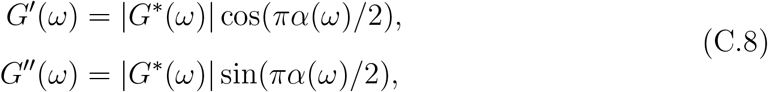

with |*G*^∗^(*ω*)| given by:

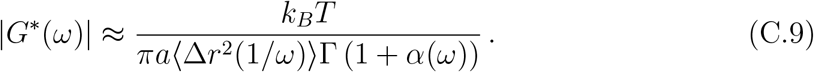

This method intrinsically assumes that the MSD will vary on logarithmic timescales, so it is expected to be more accurate when the MSD is sufficiently smooth. Moreover, the estimate of *G*^*′*^ (resp. *G*^*′′*^) becomes less accurate for *α* ≈ 1 and *α* ≈ 0. These correspond, respectively, to regimes of ballistic growth of the MSD and to a *t*-independent plateau in the MSD.

### Appendix C.2. Evans’ method

While Eq. C.9 is a straightforward technique for obtaining the frequency-dependent shear moduli, the approximate Fourier (or Laplace) transforms may affect the accuracy of the result. An improved algorithm was devised in Ref. [37] that sidesteps this by directly reconstructing a discretized complex shear modulus starting from the creep compliance *J*(*t*). The compliance is a measure of ‘deformability’ of a material, i.e. the amount of shear produced by a unit stress. For elastic solids, it is the inverse of the shear modulus:

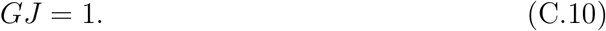

For viscoelastic materials, Eq. C.10 generalizes to a convolution in the time domain [37], which in the frequency domain again becomes simply a product:

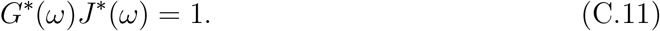

Combining Eqs. C.5 and C.11, one sees that MSD data give direct access to compliance estimates via

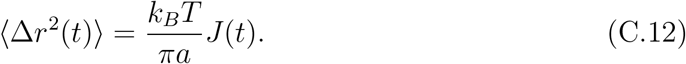

It is reasonable to assume that a viscoelastic fluid will have a long-time viscous behavior, resulting in a linear scaling *J* ∼ *t/η*, where *η* is the steady state viscosity of the material, which needs to be extrapolated from the data. As a result, the second time derivative 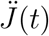 of *J*(*t*) *has* to vanish at long times. Moreover, causality requires that *J*(*t*) (and hence 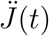) must vanish for *t <* 0 [37]. It follows that the Fourier transform of 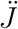 converges under all conditions, and can be used to reconstruct the Fourier transform 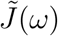 of *J*(*t*):

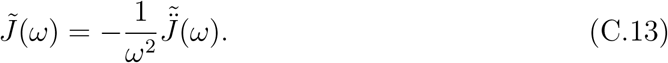

Accounting for the discontinuity at *t* = 0 induced by the time resolution of the measurements, one can then obtain

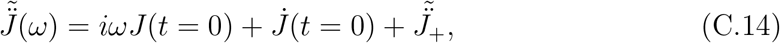

where *J*_+_ is the creep compliance restricted to strictly positive times.

In practice, we approximate *J*(*t*) with a piecewise linear function passing through the measurement points *J*_1_, …, *J*_*N*_ at sampling times *t*_1_, …, *t*_*N*_ and perform a Discrete Time Fourier Transform (DTFT) [45] on 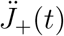 (now reduced to a series of delta functions at the measurement times) to find 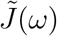 by combining equations C.11, C.13 and C.14. This leads to [37, 46] :

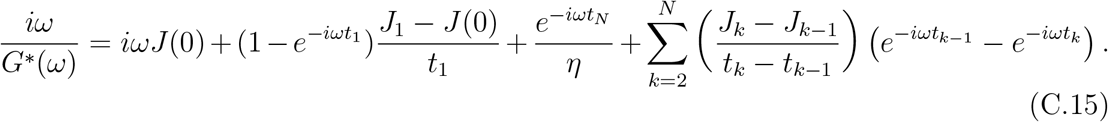

This method is independent of the sampling frequency of the *J*(*t*) measurements (which is *not* required to be uniform), but of course this sampling frequency still affects the resolution in frequency space, which is now limited by the *Nyquist–Shannon* theorem. On the other hand, the method is essentially based on the discrete implementation of an integral transform, so measurement noise effects are expected to accumulate. An improved version of the algorithm explicitly dealing with these limitations is provided in Ref. [46].

### Appendix C.3. Segment Tracking Microrheology

Above, we described the GSER for a spherical probe particle. To analyze our simulations, we in fact employ Segment Tracking Microrheology, a technique devised in Ref. [12], where filament segments are used as the probe instead of beads. This requires adapting Equations Eq. C.9 and Eq. C.13 for cylindrical probes of diameter *σ* and length *L*. This is achieved by replacing the radius of the bead *a* by an effective radius *r*_*b*_ that satisfies the following relation:

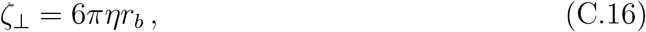

Where

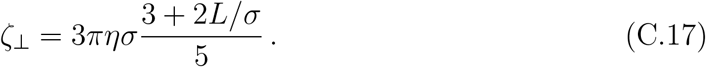

## Appendix D. Rheology simulation: Cytosim parameters corresponding to Kim et al.

In Section 3.2 of the main text, we discuss simulations performed with Cytosim aimed at studying the viscoelastic properties of actin networks. For this purpose, we prepare a cross-linked network of actin-like filaments in Cytosim, adapting parameters from Kim et al. [12]. Table D1 lists the common parameters used in the simulations. A cubic box of side length 2.8 *µ*m with periodic boundary conditions (PBC) is initialized with 500 filaments. After equilibration*‡* for 5 s, cross-linkers are added, and the system is equilibrated again for 50 s. The equilibration time with cross-linkers is chosen such that all cross-linkers are bound to two filaments, see Fig. D1 (and see Appendix D.1 below for the derivation of the analytical estimate). The timestep used is short enough to ensure that the thermal fluctuations are much smaller than the steric repulsion range of 0.007 *µm*, which prevents artifacts of filament crossing and topological inconsistencies [47].

**Figure D1:**
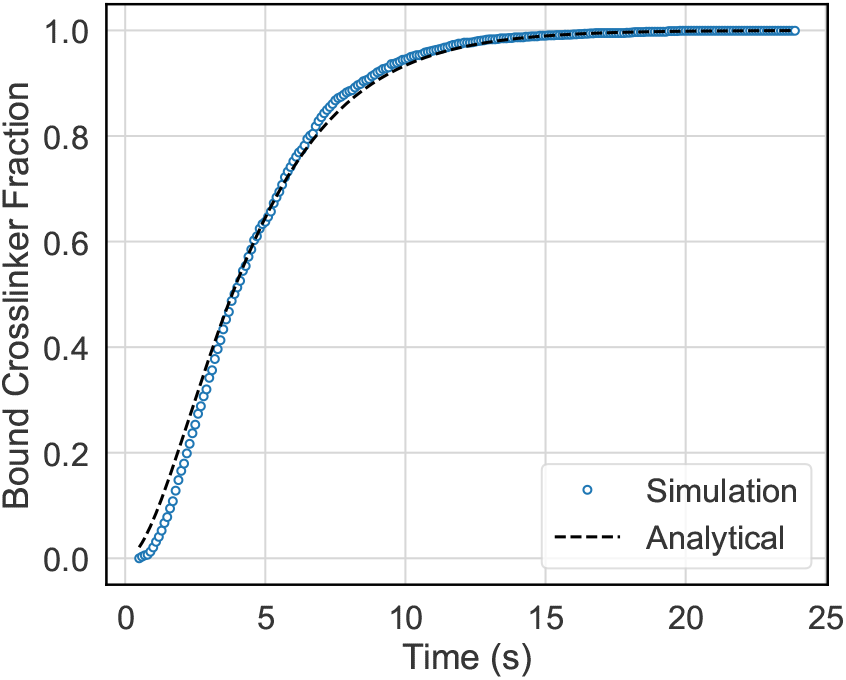
Fraction of cross-linkers bound to two filaments vs time. Derivation of the analytical expression is given in Appendix D.1

The highest meaningful frequency for measurements of *G*^∗^(*ω*) is determined by the frame-rate*§* and the lowest by the total duration of the simulation. However, when running long simulations at high frame-rates, the amount of data produced increases exponentially with each additional decade in frequency. To avoid this, different simulations are run for different ranges of frequency with a reasonable balance between duration and frame-rate. For example, we use post-equilibration measurement durations of 5 s (high frame-rate), 50 s and 500 s (longer durations), saving 500 regularly spaced frames in each case. With this setup, the data storage requirements grow linearly with the number of decades in frequency.

To obtain ensemble-averaged measurements, we ran 80 ‘trials’ for each system configuration. These were run in batches on the Göttingen University Physics Department’s GoeGrid computing cluster. With filament probes, as in the case of Segment Tracking Microrheology, random filament segments (one per filament) were tracked, with the MSD averaged over all probes across all trials.

To quantify the effect of cross-linking, we varied the number of cross-linkers in the system from *N*_*C*_ = 600 to *N*_*C*_ = 18000. The complex shear modulus was then computed from the MSD for each trial and subsequently averaged over all trials. This was found to be consistent with the complex shear modulus obtained from the ensemble-averaged MSD (Fig. 5), while simultaneously providing a measure for the variability in the shear moduli. Fig. D2 shows the curves for *G*^*′*^ and *G*^*′′*^ for the full data set of different cross-linker counts. Evans’ method was employed for this analysis to avoid low-frequency artifacts produced by Mason’s method in heavily-cross-linked networks.

**Figure D2:**
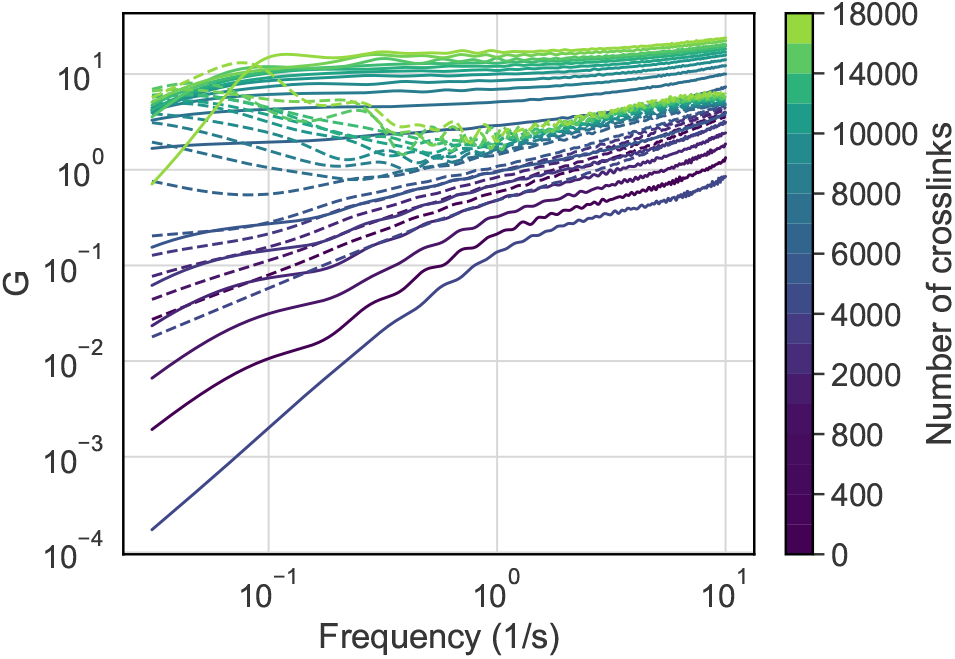
*G*^*′*^ (solid) and *G*^*′′*^ (dashed) for different cross-linker counts

**Table D1:** Parameters of benchmark simulation. Fil: filament, Cl: cross-linkers.

| Parameter | Value |
| --- | --- |
| Timestep | 0.0005 s |
| Box Length | 2.8 $\mu\text{m}$ |
| Periodicity | xyz |
| Viscosity | 0.1 Pa.s |
| Fil. length | 1.5 $\mu\text{m}$ |
| Fil. rigidity | 0.001 pN/$\mu\text{m}^2$ |
| Fil. Segmentation | 0.07 $\mu\text{m}$ |
| Fil. Steric Range | 0.007 $\mu\text{m}$ |
| Fil. Steric Force | 16.9 pN |
| Cl. Binding Rate | 10 s<sup>-1</sup> |
| Cl. Binding Range | 0.02 $\mu\text{m}$ |
| Cl. Unbinding Rate | 0 s<sup>-1</sup> |
| Cl. Stiffness | 4.23 pN/$\mu\text{m}$ |
| Cl. Diffusion | 10 $\mu\text{m}^2/\text{s}$ |

### Appendix D.1. Estimating cross-linker binding times

We can provide an analytical estimate for the equilibration time by calculating an effective binding rate for the cross-linkers and solving the corresponding Master equation. Each hand of a cross-linker binds at a rate of *κ*_*b*_ (binding rate = 10s^−1^) when within a distance *r*_*b*_ (binding range = 0.002 *µ*m) of a filament. Therefore, the effective binding rate (*k*) for a cross-linker is given by:

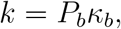

where *P*_*b*_ is the probability for a cross-linker to be in the binding range. We can estimate this using the volume fraction occupied by all *N*_*f*_ filaments, treating each as a cylinder with radius *r*_*b*_ and length *L*_*f*_ to get the effective binding rate

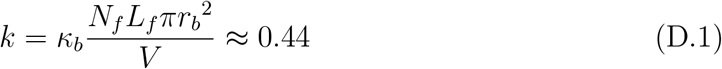

Since the network is nearly isotropic, we can assume that the binding rate is identical for both free and singly-bound cross-linkers. Using this, we can write down the Master equation for this system, where *ϕ*_*f*_, *ϕ*_*s*_ and *ϕ*_*d*_ are the fractions of free, single-bound and double-bound cross-linkers, respectively:

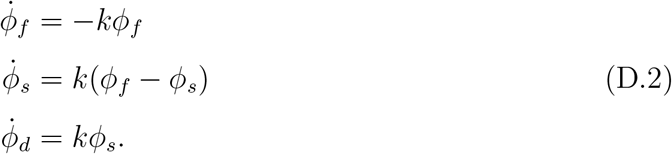

Solving this Master equation with an initial condition *ϕ*_*f*_ (0) = 1 we obtain an analytical expression for the fraction of bound cross-linkers:

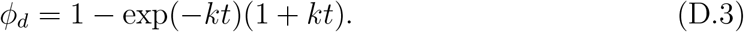

We find that this analytical approximation is in good agreement with the fraction of bound cross-linkers measured during the equilibration phase of the simulation (Fig. D1). Importantly, we note that Eq. D.3 is independent of the total number of cross-linkers, permitting the use of a constant equilibration time for simulations with different cross-linker counts.

## Appendix E. Scaling argument for *ω*^3*/*4^ power-law of *G*^*′′*^

Fig. 3 shows that, for frequencies above the crossing point, the loss modulus of the cross-linked actin network obtained using Mason’s method scales as *ω*^3*/*4^. The result can be explained on the basis of the high frequency response of semiflexible, entangled polymer networks, as extensively explained in [16] and [38, 48]. For completeness we provide a brief qualitative picture here, to highlight the physical assumptions underlying the observed scalings.

When strained at high frequency, the network response is dominated by the single polymer’s transversal oscillations, while the cross-linkers impede filament tangential sliding. By simple scaling arguments, oscillations must increase with temperature *T* and with the wavelength *λ* of the bending mode, while decreasing with the bending modulus *κ*. Defining *u* as the transversal displacement field from the straight filament configuration, we then expect typical amplitudes 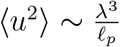, where ℓ_p_is the persistence length (ℓ_*p*_ ∼ *κ/T* ). It is well known that transversal fluctuations for a polymer in a viscous solution have a dispersion relation *ω* ∼ *λ*^−4^ [49], so we expect the typical bending mode relaxing after time *t* to be of wavelength *λ* ∼ *t*^1*/*4^. Knowing that the transversal fluctuations are dominated by the longest unconstrained bending mode [16] (of wavelength of the order of the entanglement length ℓ_*e*_ in our case), then *typical* end-to-end polymer length fluctuations will scale as ⟨*u*^2^⟩ ∼ *t*^3*/*4^, implying a shear modulus ∼ (*iω*)^3*/*4^ [13, 32]. The derivation of this high frequency scaling thus relies on the assumption that the cross-linked network of semiflexible polymers has ℓ_*e*_ *<* ℓ_*p*_; in the other case, which is not explored numerically in this work, simple fluctuations of the order ∼ *t*^1*/*2^ are expected, as can be easily seen from the above argument considering the longest unconstrained bending mode to be of length ℓ_*p*_.

## Appendix F. AI documentation

During the preparation of this work the authors used chatGPT models 4o and 5 in order to improve manuscript writing quality. After using this tool, the authors reviewed and edited the content as needed and take full responsibility for the content of the publication.

## Footnotes

‡ Filaments are initially placed randomly without any bending. This short equilibration imparts Brownian fluctuations to the segments.

§ This is the inverse of the time between two saved frames (*t*_*f*_ ), which is often longer than the *timestep*.

## Notes

https://hdl.handle.net/21.11101/0000-0007-FE56-B

https://doi.org/10.25625/JGDI8Y

### Competing Interest Statement

The authors have declared no competing interest.

